# Morphological variation and phylogeny of *Karenia selliformis* (Gymnodiniales, Dinophyceae) in an intensive cold-water algal bloom in eastern Hokkaido, Japan in September–November 2021

**DOI:** 10.1101/2021.12.13.472515

**Authors:** Mitsunori Iwataki, Wai Mun Lum, Koyo Kuwata, Kazuya Takahashi, Daichi Arima, Takanori Kuribayashi, Yuki Kosaka, Natsuki Hasegawa, Tsuyoshi Watanabe, Tomoyuki Shikata, Tomonori Isada, Tatiana Yu. Orlova, Setsuko Sakamoto

**Affiliations:** Graduate School of Agricultural and Life Sciences, University of Tokyo, Tokyo 113-8657, Japan; Central Fisheries Research Institute, Hokkaido Research Organization, Yoichi, Hokkaido 046-8555, Japan; Fisheries Research Institute, Aomori Prefectural Industrial Technology Research Center, Hiranai, Aomori 039-3381, Japan; Fisheries Resources Institute, Japan Fisheries Research and Education Agency, Kushiro, Hokkaido 085-0802, Japan; Fisheries Technology Institute, Japan Fisheries Research and Education Agency, Goto, Nagasaki 853-0508, Japan; Akkeshi Marine Station, Field Science Center for Northern Biosphere, Hokkaido University, Akkeshi, Hokkaido 088-1113, Japan; National Scientific Center of Marine Biology, Far East Branch, Russian Academy of Sciences, Vladivostok 690041, Russia; Fisheries Technology Institute, Japan Fisheries Research and Education Agency, Hatsukaichi, Hiroshima 739-0452, Japan

**Keywords:** cold-water bloom, dinoflagellates, harmful algal blooms, Karenia, Karlodinium, Takayama

## Abstract

Harmful algal blooms responsible for mass mortalities of marine organisms have so far been rare in Hokkaido, northern Japan, although fish killing blooms have been frequently reported from western Japanese coasts. In September–November 2021, a huge and prolonged cold-water bloom occurred along the Pacific coast of eastern Hokkaido, Japan, and was associated with intensive mortalities of sea urchin, fish, octopus, shellfish, etc. In this study, morphology and phylogeny of the dominant and co-occurred unarmored dinoflagellates of the Kareniaceae in the bloom were examined by using light microscopy, scanning electron microscopy and molecular phylogeny inferred from ITS and LSU rDNA (D1–D3) sequences. Morphological observation and molecular phylogeny showed that the dominant species was *Karenia selliformis*, with co-occurrences of other kareniacean dinoflagellates, *Kr. longicanalis*, *Kr. mikimotoi*, *Karlodinium* sp., *Takayama* cf. *acrotrocha*, *Takayama tuberculata* and *Takayama* sp. The typical cell forms of *K. selliformis* in the bloom were discoid, dorsoventrally flattened, and larger than the cell sizes in previous reports, 35.3–43.6 (39.4±2.1) μm in length. Transparent cells of *Kr. selliformis* lacking or having several shrunken chloroplasts and oil droplets were also found. Cells of *Kr. selliformis* had morphological variation, but the species could be distinguished from other co-occurred *Karenia* species by its numerous (46–105) and small granular (2.9–4.6 μm in diameter) chloroplasts and the nucleus positioned in the hypocone. Cell density of *Kr. selliformis* exceeding 100 cells/mL was recorded in the range of temperature 9.8–17.6°C. The rDNA sequences determined from *Kr. selliformis* in the blooms of Hokkaido, Japan in 2021 were identical to those from another bloom in Kamchatka, Russia in 2020.

**Highlights:**

- A marine fauna-destructive harmful algal bloom in the Pacific coast of eastern Hokkaido, Japan in September–November 2021 was dominated by *Karenia selliformis*.
- Cells of *Karenia selliformis* typical in the bloom were discoid and possessing numerous small chloroplasts, approximately 70 in number.
- Cells of *Karenia selliformis* showed morphological variation in size and shape, and transparent motile cells lacking or having degraded chloroplasts were also present.
- Co-occurred kareniaceans in the bloom were *Karenia longicanalis*, *Karenia mikimotoi*, *Karlodinium* sp. and *Takayama* spp.
- rDNA sequences of *Karenia selliformis* in the blooms of Hokkaido in 2021 and Kamchatka in 2020 were identical, which belong to the group I of *Kr. selliformis*.

## 1. Introduction

The unarmored dinoflagellates in the Kareniaceae are responsible for mass mortalities of fish and other organisms in coastal marine environments (Lundholm et al., 2009 onwards). Among the genera of this family, *Asterodinium* Sournia, *Brachidinium* Sournia, *Gertia* K.Takahashi et Iwataki, *Karenia* G.Hansen et Moestrup, *Karlodinium* Larsen, *Shimiella* Ok, Jeong, Lee et Noh, and *Takayama* de Salas, Bolch et Hallegraeff, species in *Karenia*, *Karlodinium* and *Takayama* have been reported as harmful algal bloom (HAB) causative species (Daugbjerg et al., 2000; de Salas et al., 2003; Benico et al., 2019; Takahashi et al., 2019; Ok et al., 2021). In the Northwest Pacific, fisheries damage has been reported with kareniacean blooms of *Karenia mikimotoi* (Miyake et Kominami ex Oda) G.Hansen et Moestrup, *Karenia longicanalis* Yang, Hodgkiss et G.Hansen, *Karenia papilionacea* Haywood et Steidinger, *Karlodinium digitatum* (Yang, Takayama, Matsuoka et Hodgkiss) Gu, Chan et Lu, *Karlodinium australe* de Salas, Bolch et Hallegraeff, *Karlodinium veneficum* (Ballantine) Larsen, and *Takayama acrotrocha* Larsen, both in the Southeast and East Asia (Yang et al., 2000, 2001; Lim et al., 2014; Tang et al., 2014; Yamaguchi et al., 2016; Li et al., 2019; Benico et al., 2020). In Japanese coastal waters, *Kr. mikimotoi* is one of the most noxious harmful microalgae, which has caused severe mortalities of farmed fish and shellfishes (Oda, 1935; Okaichi, 1987, 2003; Imai et al., 2006; Aoki et al., 2020; Sakamoto et al., 2021). Other kareniacean dinoflagellates such as *Kr. longicanalis*, *Kr. papilionacea* and *Kl. digitatum* have also been observed (e.g., Yang et al., 2000; Yamaguchi et al., 2016), as well as *Kr. mikimotoi*. These species have proliferated mainly in the coasts of western Japan, such as Seto Inland Sea and Kyushu area (Okaichi, 1987; Imai et al., 2006; Sakamoto et al., 2021).

In the coastal waters of Hokkaido, northeast Japan, harmful microalgae noxious for marine organisms have rarely been observed, although the paralytic shellfish toxin producer *Alexandrium catenella* (Group I) has commonly occurred (Shimada et al., 2016a; Natsuike et al., 2021). Recently, a bloom of *Kr. mikimotoi* was first found in Hakodate Bay, southern Hokkaido in 2015, while the negative impact was not intensive, and cells of harmful raphidophyte *Chattonella marina* (Subrahmanyan) Hara et Chihara and dinoflagellate *Cochlodinium polykrikoides* Margalef (= *Margalefidinium polykrikoides*) were detected during summer (Shimada et al., 2016a, 2016b; Kakumu et al., 2018). Blooms of these harmful microalgae are common in the coasts of western Japan, and even in Southeast Asia (Imai et al., 2006; Yamatogi et al., 2006; Iwataki et al., 2008, 2015; Imai and Yamaguchi, 2012; Furuya et al., 2018; Thoha et al., 2019; Lum et al., 2021; Sakamoto et al., 2021; Yñiguez et al., 2021). It was assumed that the motile cells of these harmful species recently detected in the coastal Hokkaido might be transported by the Tsushima Warm Current from the Sea of Japan (Shimada et al., 2016a; Sildever et al., 2019).

In September–November 2021, a massive dinoflagellate bloom abruptly occurred along the Pacific coast of eastern Hokkaido (Kuroda et al., 2021). During the bloom, intensive mortalities of marine organisms, such as sea urchins (*Strongylocentrotus intermedius* and *Mesocentrotus nudus*), wild salmon (*Oncorhynchus keta*) captured by stationary net-traps, octopus (*Paroctopus conispadiceus*), whelks (*Neptunea* spp.), chitons (*Cryptochiton stelleri*), bivalves (e.g., *Pseudocardium sachalinense*), etc., were observed (e.g., Misaka and Ando, 2021), although the killing mechanisms and causative compounds are not elucidated yet. Mass mortality of gray sea urchin and cucumaria was observed also in Kunashiri and Habomai Islands since 1st October 2021 (Sergei Maslennikov, pers. commun.). The greenish brown algal bloom with yellowish form was first recognized on 13–16th September 2021, due to the seawater discoloration around the pier of Akkeshi Marine Station, Hokkaido University, and mortalities of cultured marine organisms in Japan Fisheries Research and Education Agency, Kushiro, Hokkaido. The bloom was subsequently expanded along the Pacific coast of eastern Hokkaido, in the cold-water mass (Kuroda et al., 2021). Although the negative impacts including benthic organisms were not fully realized, the provisional estimation of fisheries damage might exceed the largest record in Japan, i.e., 7.1 M USD due to *Chattonella* bloom in the Seto Inland Sea in 1972 (Okaichi, 1987; Sakamoto et al., 2021).

The aim of this study was an unambiguous identification of the bloom causative species as a first step to understand the trait of this algal bloom, such as blooming mechanism, toxicity, and negative impacts on marine fauna. For this purpose, the dominant and associated dinoflagellates in the Hokkaido bloom 2021 were isolated and used for single cell PCR, culture establishment, and light and scanning electron microscopy. Morphology and phylogeny showed the causative species was *Karenia selliformis* Haywood, Steidinger et MacKenzie, which has no previous bloom record in Japan.

## 2. Materials and Methods

### 2.1. Samples of Hokkaido bloom 2021

The satellite ocean color images of chlorophyll *a* concentration and sea surface temperature in Hokkaido, on 9th October, derived from the Second generation GLobal Imager (SGLI) aboard the Global Change Observation Mission-Climate (GCOM-C) launched by Japan Aerospace Exploration Agency (JAXA), were obtained from the JASMES website (https://www.eorc.jaxa.jp/JASMES/SGLI_STD/daily.html?area=j&prod=CHLA&drct=D) (Fig. 1). Measured seawater temperatures were obtained on 9th October, except for those in Taiki and Toyokoro measured on 6th October (Fig. 1B). For identification of HAB causative species, motile cells were isolated from the bloom samples collected from the coasts of eastern Hokkaido, after one to two days of transportation, and used for single cell PCR and culture establishment (Fig. 1, Table 1). Cells were counted to estimate cell densities in the Central Fisheries Research Institute, Hokkaido Research Organization. For morphological and phylogenetic comparison of kareniaceans found in Hokkaido bloom, cultures of *Kr. selliformis* and *Kr. longicanalis* previously established from Aomori, Japan were also used (Table 1).

**Table 1.** Samples examined in this study. Cultures and single cell isolates of unarmored dinoflagellates in the Kareniaceae obtained from the bloom of eastern Hokkaido 2021, and other cultures for comparison.

| Date | Location | Coordinates | Temperature | Salinity | Specimen |
| --- | --- | --- | --- | --- | --- |
| 20 Aug<br>2019 | Hiranai,<br>Aomori | 40.946194°N<br>140.861125°E | 23.0°C | 34.0 | Culture of *Kr. selliformis* (MoKr600) |
| 23 Sept<br>2020 | Noheji,<br>Aomori | 40.916667°N,<br>141.116667°E | 22.0°C | 33.0 | Culture of *Kr. longicanalis* (AmKr650) |
| 20 Sept<br>2021 | Akkeshi,<br>Hokkaido | 43.021448°N,<br>144.836646°E | 17.1°C | 31.9 | Cultures of *Kr. longicanalis*<br>(LAkKL297, LAkKL298, LAkKL299,<br>LAkKL300), *Takayama* sp.<br>(LAkTak296) |
| 20 Sept<br>2021 | Katsurakoi<br>Port, Kushiro,<br>Hokkaido | 42.944228°N<br>144.447845°E | 17.8°C | 31.9 | Culture of *T. cf. acrotracha* (KKsTk67) |
| 24 Sept<br>2021 | Katsurakoi<br>Port, Kushiro,<br>Hokkaido | 42.944228°N<br>144.447845°E | 16.5°C | 28.2 | Cultures of *Kr. selliformis* (KKsKs74,<br>KKsKs75), *Kr. mikimotoi* (KKsKm70,<br>KKsKm71, KKsKm72) |
| 25 Sept<br>2021 | Katsurakoi<br>Port, Kushiro,<br>Hokkaido | 42.944228°N<br>144.447845°E | 13.2°C | 30.0 | Isolates of *Kr. selliformis* (K1, K3, K7,<br>M2, M3, M4, M5, M6, M7, M8),<br>cultures of *Kr. selliformis* (LK2Ks306,<br>LK2Ks307) |
| 29 Sept<br>2021 | Katsurakoi<br>Port, Kushiro,<br>Hokkaido | 42.944387°N,<br>144.445859°E | 16.2°C | 32.3 | Cultures of *Kr. mikimotoi* (KKsKm73),<br>*Takayama* sp. (KKsTk69,<br>LK929Tak330) |
| 13 Oct<br>2021 | Asahihama<br>Port, Taiki,<br>Hokkaido | 42.420317°N,<br>143.392300°E | 14.4°C | 30.5 | Isolates of *Kr. selliformis* (AsahiL2,<br>AsahiL7) |
| 14 Oct<br>2021 | Otsu Port,<br>Toyokoro,<br>Hokkaido | 42.675300°N,<br>143.640867°E | 13.6°C | 27.9 | Isolates of *Kr. selliformis* (OT2, OT3,<br>OT4), |

**Fig. 1.**
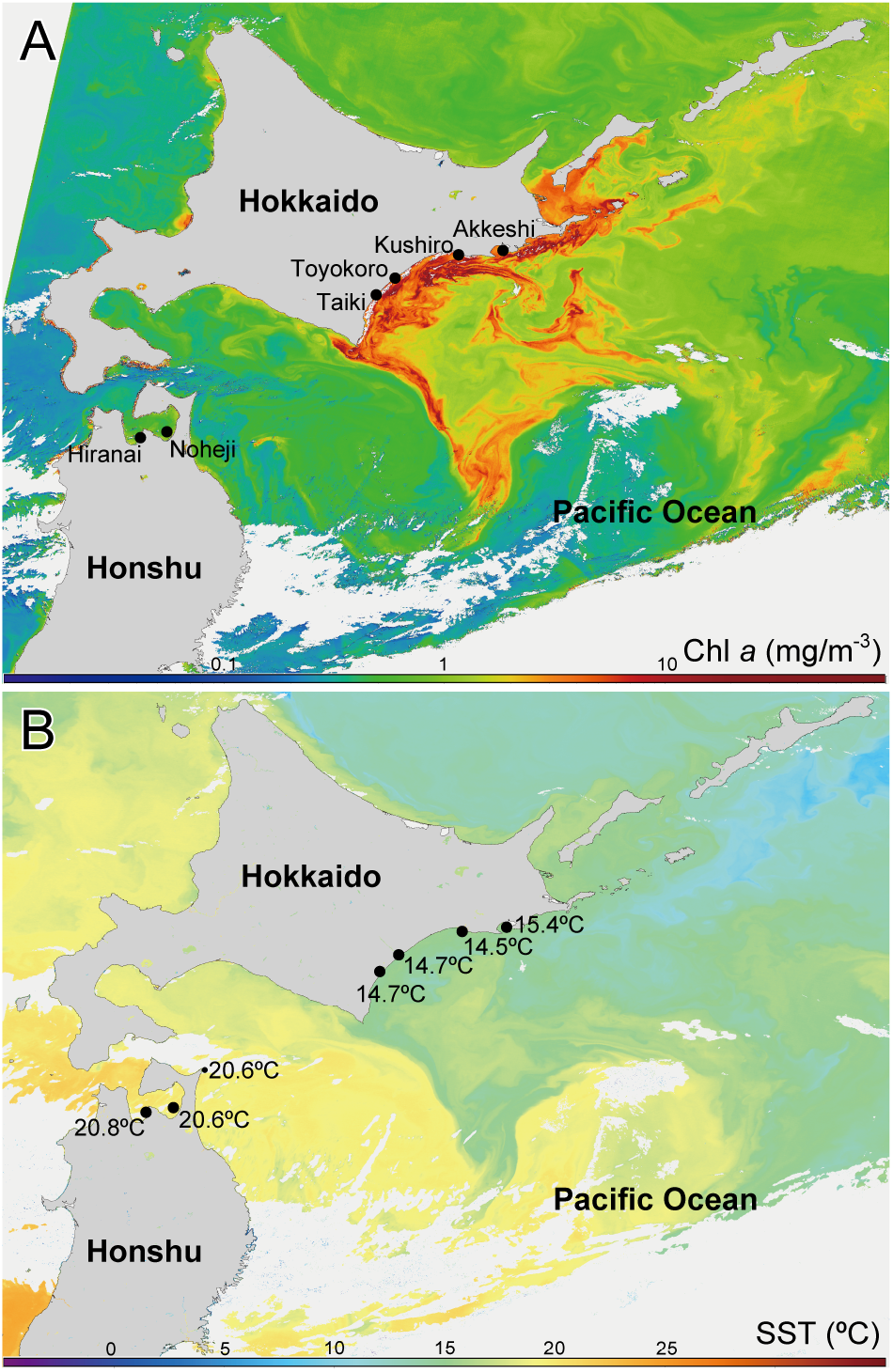
Satellite ocean color images of chlorophyll *a* (A) and sea surface temperature (B) of Hokkaido, Japan on 9th October 2021. Measured seawater temperatures were obtained on 9th October, except for those in Taiki and Toyokoro measured on 6th October. Satellite images were obtained from the Second generation GLobal Imager (SGLI) aboard the Global Change Observation Mission-Climate (GCOM-C) of Japan Aerospace Exploration Agency (JAXA). Black circles indicate sampling locations in this study.

### 2.2. Cultures and light microscopy

Cells were isolated by capillary pipetting under inverted microscopes Olympus CKX41 and CKX53 (Olympus, Tokyo, Japan), and cultures were maintained in 1/2 IMK (Wako, Tokyo, Japan) or modified K/2 medium, under the conditions of 18–20°C, 30 psu, and a 12:12 h light:dark cycle with 40–50 μmol photons m^−2^ s^−1^ light illumination. Isolated single cells were also used for DNA amplification and molecular phylogeny (Table 1).

Cells were observed using a light microscope Zeiss Axioskop 2 (Carl Zeiss, Gӧttingen, Germany) equipped with a digital camera Axiocam 305 color (Carl Zeiss), and an inverted microscope Olympus IX71 with a digital camera DP73 (Olympus, Tokyo, Japan). Autofluorescence of chloroplast and nucleus stained by DAPI were observed under a UV excitation (Zeiss filter set 02), according to previous study (Takahashi et al., 2014).

### 2.3. Scanning electron microscopy (SEM)

For SEM observation, cell fixations were slightly modified from Lum et al. (2019). Cells were fixed with 2% (w/v) OsO_4_ for 30 min on a poly-L-lysine-coated glass plate at room temperature and were rinsed twice with distilled water for 30 min each. Cells were dehydrated through an ethanol series of 10, 30, 50, 75, 90, and 95% (15 min each) and twice in 100% ethanol (30 min each), which was then replaced with isoamyl acetate (15 min). Cells were dried using a JCPD-5 critical point dryer (JEOL, Tokyo, Japan) and sputter-coated with platinum-palladium. They were then observed using an S-4800 SEM (JEOL, Tokyo, Japan) operated at 1.0–5.0 kV.

### 2.4. DNA sequencing and phylogenetic analyses

For rDNA sequencing, previously described primers were used for the amplification and sequencing of ITS region and LSU rDNA (Kawami et al., 2006; Iwataki et al., 2007, 2008; Takahashi et al., 2015). PCR were conducted by the following processes. Cells in seawater samples and culture strains were picked up by capillary pipetting, and were put into sterilized distilled water. After observation under an inverted microscope Olympus IX71, cells were transferred into PCR strips to use directly as a PCR template. The initial PCR was conducted by use of a forward primer SR10 (5’-AGG TCT GTG ATG CCC TTA GA-3’) and a reverse primer 28-1483R (5’-GCT ACT ACC ACC AAG ATC TGC-3’). By using the amplicons as a template, the nested PCR was conducted with a forward primer ITSA (5’-GTA ACA AGG THT CCG TAG GT-3’) and a reverse primer D3B (5’-TCG GAG GGA ACC AGC TAC TA-3’). PCR was conducted by use of Ex Taq polymerase (Takara, Shiga, Japan) at a reaction volume of 10 μL, according to the manufacturer’s protocol. The thermal cycling conditions included an initial denaturation at 94°C (1 min), followed by 35 cycles of the following 3 steps: 94°C (20 sec), 50°C (30 sec), and 72°C (2.5–3.5 min) and finally, an elongation step of 72°C (7 min). Amplicons were purified by using a QIAquick PCR Purification Kit (Qiagen Genomics, Bothell, WA, USA), following the manufacturer’s protocol. Amplicons were sequenced by Eurofins Genomics Inc. (Tokyo, Japan).

Determined ITS and LSU rDNA sequences were aligned with sequences obtained from GenBank by using Clustal X (Thompson et al., 1997) and obviously misaligned sites were then manually corrected. To ease the following analytical process, identical sequences were omitted or concatenated into a single sequence; but sequences determined in the present study were not concatenated. Phylogenetic trees for each marker were constructed by Bayesian inference (BI), using a MrBayes v.3.1.2 software (Ronquist and Huelsenbeck, 2003). The best substitution models selected by a MrModeltest v.2.3 (Nylander, 2008) were general time reversible (GTR) plus gamma distribution (G = 0.3907) for ITS, and GTR plus gamma (G = 0.4421) plus the proportion of invariable sites (I = 0.3336) for LSU rDNA. Posterior probabilities (PP) of BI were calculated using 500,000 Markov chain Monte Carlo generations with four chains and trees were sampled every 100 generations. For a confirmation for convergence of the chains, the average standard deviations of the split frequencies after calculations were below 0.01 (0.004093 for ITS, 0.009149 for LSU rDNA). Outgroups selected were *Karlodinium armiger* Bergholtz, Daugbjerg et Moestrup, *Kl. australe* de Salas, Bolch et Hallegraeff and *Kl. decipiens* de Salas et Laza-Martinez for ITS, and *Gyrodinium dominans* Hulbert for LSU rDNA. For ITS and LSU rDNA trees, PP supports were also estimated by 500 replicates of Bootstrap support (BS) values of maximum likelihood (ML) phylogeny by MEGA 6.0 (Tamura et al., 2013).

## 3. Results

### 3.1. Seawater temperature of the Hokkaido bloom 2021

Cell densities of *Kr. selliformis* were counted in the seawater samples collected from eastern Hokkaido, from 22nd September to 25th November 2021. The temperature ranges of which *Kr. selliformis* cells occurred were 8.1–19.0°C, cell densities of >100 cells mL^−1^ were 9.8–17.6°C, and >1,000 cells mL^−1^ were 11.0–17.3°C. On 27th October, high cell densities of 10,560 cells mL^−1^ at Toyokoro and 9,600 cells mL^−1^ at Hiroo were found under 11.9°C and 11.3°C, respectively. At the end of November, seawater temperature decreased to approximately 9°C or below, and the cell densities have also decreased to <10 cells mL^−1^. Cells of *Kr. selliformis* were not detected from the coasts of Aomori, Honshu, in September–November 2021. Detailed environmental condition of the *Kr. selliformis* bloom will be reported elsewhere after the analyses will be carried out.

### 3.2. *Morphology of* Karenia selliformis *in the bloom 2021*

Observed bloom samples were dominated by *Kr. selliformis*, and other unarmored dinoflagellates of the Kareniaceae were scarcely found. Cells of *Kr. selliformis* had variations in size and shape, and they could be distinguished from other kareniaceans based on their numerous and small-sized chloroplasts. Transparent cells of *K. selliformis* were identified with the help of molecular identification (see below).

Cells of *Kr. selliformis*, the typical form in the bloom, were discoid and dorsoventrally flattened, 35.3–43.6 μm (mean 39.4±2.1 μm, *n* = 40) long and 31.8–44.7 μm (mean 37.3±3.2 μm, *n* = 40) wide (Figs 2, S1, Table 2). In culture, cells were slightly ellipsoidal, 38.6–48.8 μm (mean 43.0±2.6 μm, *n* = 40) long and 30.9–40.1 μm (mean 36.3±2.5 μm, *n* = 40) wide. The roundish conical epicone and the hemispherical hypocone were similar in size, and the antapical depression was present (Fig. 2A–C). The ventral side was flat and smooth, while the dorsal was bulged and wrinkled with longitudinal striations on the surface (Fig. 2D–F). The cingulum was positioned at median and declining 2–3 times of its own width (Fig. 2A). The sulcus was deep and reached to the antapical depression, and the anterior sulcal extension was short and wide with an open end. The apical structure complex (ASC, = apical groove) was straight, extended from the right side of the sulcal extension toward the dorsal, and terminated at one third of the epicone (Fig. 2A, D–F). Chloroplasts were granular or strap-shaped, irregularly curved with an internal pyrenoid, situated in the periphery of the cell, and numerous in the typical cell (Fig. 2B, G, H). The nucleus was positioned in the hypocone, laterally elongated, and the left side was slightly larger and extended to the anterior (Fig. 2C, G, H). Two pusules, each positioned near the longitudinal and transverse flagellar canals, were present (Fig. 2B). Secretion of irregular-shaped translucent material and ejection of trichocysts could be seen (Fig. 2I). In the bloom samples, most cells formed the typical shape and size mentioned above, and morphological variations such as smaller and ellipsoidal cells were also observed (Figs 1J–L, S2). These smaller cells possessed the spherical or elliptical nucleus located in the hypocone, and chloroplasts similar in size to those of the typical cells.

**Table 2.** Morphological comparison of *Karenia longicanalis*, *Kr. mikimotoi* and *Kr. selliformis* observed in the Hokkaido bloom and previous reports.

|  | *Kr. longicanalis* | *Kr. longicanalis* | *Kr. longicanalis*<br>(as *K. umbella*) | *Kr. mikimotoi* | *Kr. mikimotoi* | *Kr. selliformis* | *Kr. selliformis* | *Kr. selliformis* |
| --- | --- | --- | --- | --- | --- | --- | --- | --- |
| Cell shape | Ovoid |  | Ovate, slightly dorsoventrally flattened | Conical epicone, hemispherical hypocone, dorsoventrally flattened | Conical or hemispherical epicone, hemispherical hypocone | Discoidal to ellipsoidal, dorsoventrally flattened | Oval in shape or longer than wide |  |
| Length<br>(mean±s.d.) | 28.6–37.6 µm<br>(32.9±2.2 µm) | 17.5–35.0 µm<br>(25.9±5.5 µm) | 29–42 µm<br>(35.9±3.4 µm) | 24.6–35.1 µm<br>(31.2±3.0 µm) | 23.9–37.7 µm<br>(32.8±3.4 µm) | 35.3–43.6 µm<br>(39.4±2.1 µm) | 20–27 µm<br>(23.6±0.2 µm) | 25–30 µm |
| Width<br>(mean±s.d.) | 23.4–30.7 µm<br>(26.5±2.0 µm) | 10.0–22.5 µm<br>(21.1±4.3 µm) | 21–32 µm<br>(26.6±2.9 µm) | 21.9–30.9 µm<br>(26.8±2.4 µm) | 21.6–36.4 µm<br>(30.6±3.8 µm) | 31.8–44.7 µm<br>(37.3±3.2 µm) | 16–24 µm<br>(18.5±0.3 µm) | 20–25 µm |
| Nucleus | Ellipsoid, central or slightly posterior | Round and located in the central, surrounded by a nuclear capsule | Round to ellipsoidal, central | Round in the left hypocone, or ellipsoid in the left of the cell | Reniform or pyriform situated in the left part of the cell | Reniform, laterally elongated, in the hypocone | Oval, elliptical, elongated reniform in the horizontal plane in the hypocone | Oblong and occupied most of the hyposome in a horizontal position |
| Chloroplast shape | Multilobed with a central pyrenoid | Round | Irregular, shallow, multilobed and strap-shaped | Bean-shaped with a lateral pyrenoid | Elongated, irregular | Subspherical to strap-shaped, with a pyrenoid | Reniform |  |
| Chloroplast number | 4–7 (6.1±0.6) | About 30 | Approx. 20 | 13–39<br>(24.5±6.4) | Generally 10–15 or more | 46–105<br>(70.6±15.8) | Several | Several |
| Chloroplast size<br>(major axis) | 9.0–19.4 µm<br>(14.5±2.4 µm) |  |  | 4.7–10.6 µm<br>(7.3±1.5 µm) |  | 2.9–4.6 µm<br>(3.7±0.4 µm)<br>(subspherical)<br>6.2–15.1 µm<br>(9.4±1.9 µm)<br>(strap-shaped) |  |  |
| References | This study | Yang et al. (2001) | de Salas et al. (2004) | This study | Hansen et al. (2000) | This study | Haywood et al. (2004) | Mardones et al. (2020) |

**Fig. 2.**
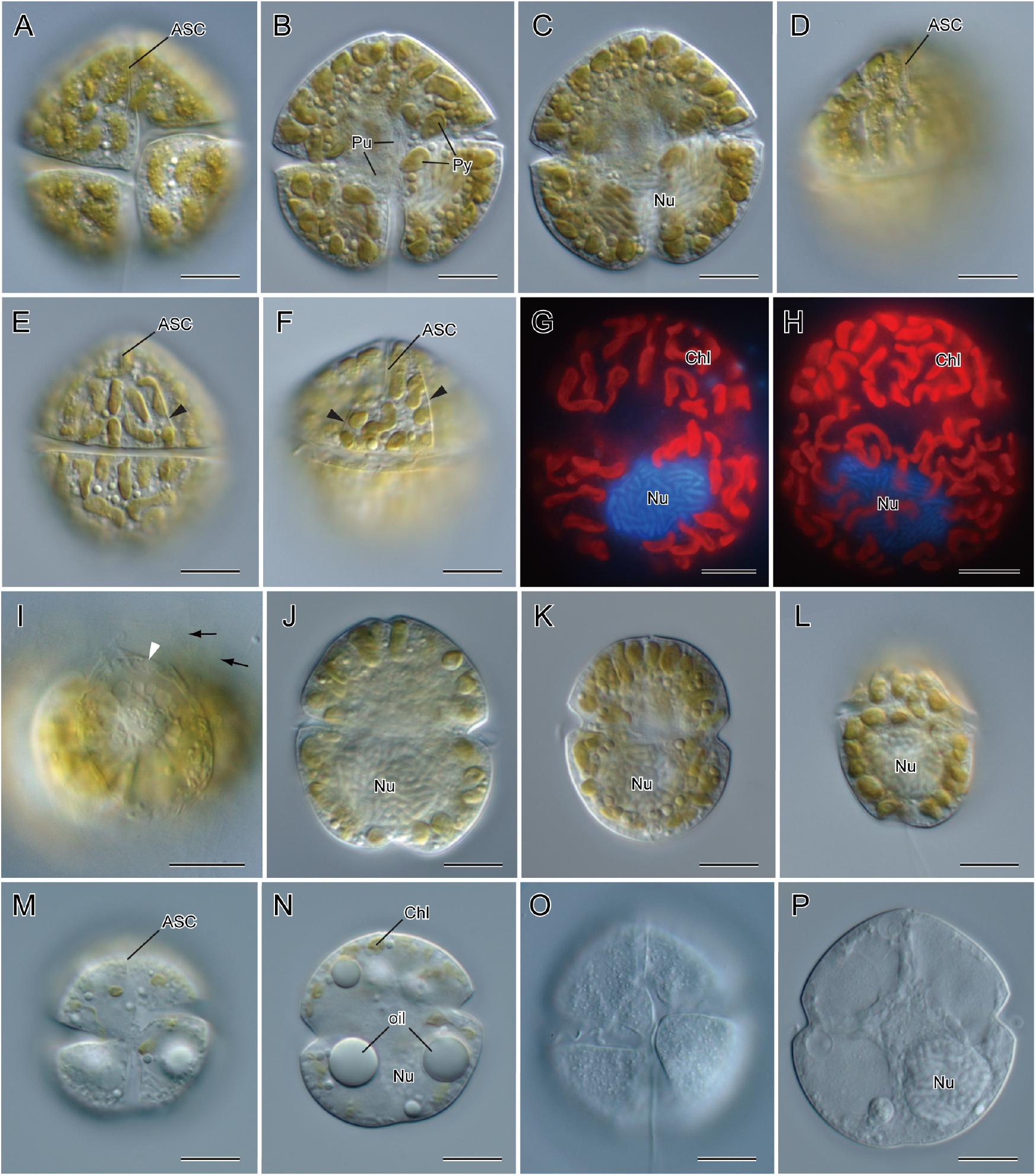
Morphological variations of *Karenia selliformis* found in the bloom 2021, light and fluorescence microscopy. (A–C) Typical cell shape in the bloom, from ventral, showing the straight apical structure complex (ASC, = apical groove), pusules (Pu), pyrenoids (Py) in chloroplasts, and nucleus (Nu) posteriorly located. (D–F) Typical cell shape, from apico-dorsal and dorsal, note the ASC and bulging dorsal side with striations (arrowheads). (G, H) Fluorescence microscopy showing many chloroplasts (Chl) and DAPI-stained nucleus (Nu). (I) Secreted mucilaginous material (white arrowhead) and released trichocysts (arrows), antapical view. (J–L) Size and morphological variations of the cells, note the nucleus (Nu) positioned posteriorly. (M, N) A transparent cell with the straight apical structure complex (ASC), small and irregular-shaped chloroplasts (Chl), oil droplets (oil), and nucleus (Nu). (O, P) A transparent cell without visible chloroplasts, with nucleus (Nu). Scale bars = 10 μm.

Transparent motile cells, having several shrunken and irregular-shaped chloroplasts, or lacking chloroplasts, also occurred in the bloom samples (Fig. 2M–P). These reduced chloroplasts still showed the autofluorescence, and were not observed in the cells without chloroplast (not shown). They were smaller than the typical form, more or less spherical, and the straight ASC could be observed (Fig. 2M, O). Several oil droplets, which were apparently larger than those found in the typical cell form, were conspicuous and common in the transparent cells (Figs 1M, N, S2). The nucleus was spherical and positioned in the hypocone (Fig. 2N, P). These transparent cells, even when lacking all chloroplasts, were actively swimming with a longitudinal and a transverse flagellum (Fig. 2O). Transparent cells similar to the typical cell form were rarely observed (Fig. 2O, P).

Scanning electron microscopy showed the cell surface of *Kr. selliformis* with polygonal amphiesmal vesicles (Fig. 3). The cingulum was positioned in the middle and descending about the cingular width (Fig. 3A, B). The anterior ridge of the cingulum was clearly seen as a line, while the posterior margin was not apparent (Fig. 3A–C). The right side of the sulcus, at the junction of cingular end, was slightly bulged (Fig. 3A, B). The anterior sulcal extension was shallow and the distal end was open (Fig. 3A, B). The straight apical structure complex (ASC, = apical groove) consisted of ridged knob vesicles at the left side, smooth vesicles in the furrow, and a ridge without knobs at the right side (Fig. 3B). The ASC started near the right side of the sulcal extension, extended toward the cell apex, and terminated at the upper part of the epicone in the dorsal side. The surface was rugose in the dorsal, which was consistent with LM observation (Fig. 3C). The ventral pore was absent, and the lateral pore was not observed (Fig. 3A–C).

**Fig. 3.**
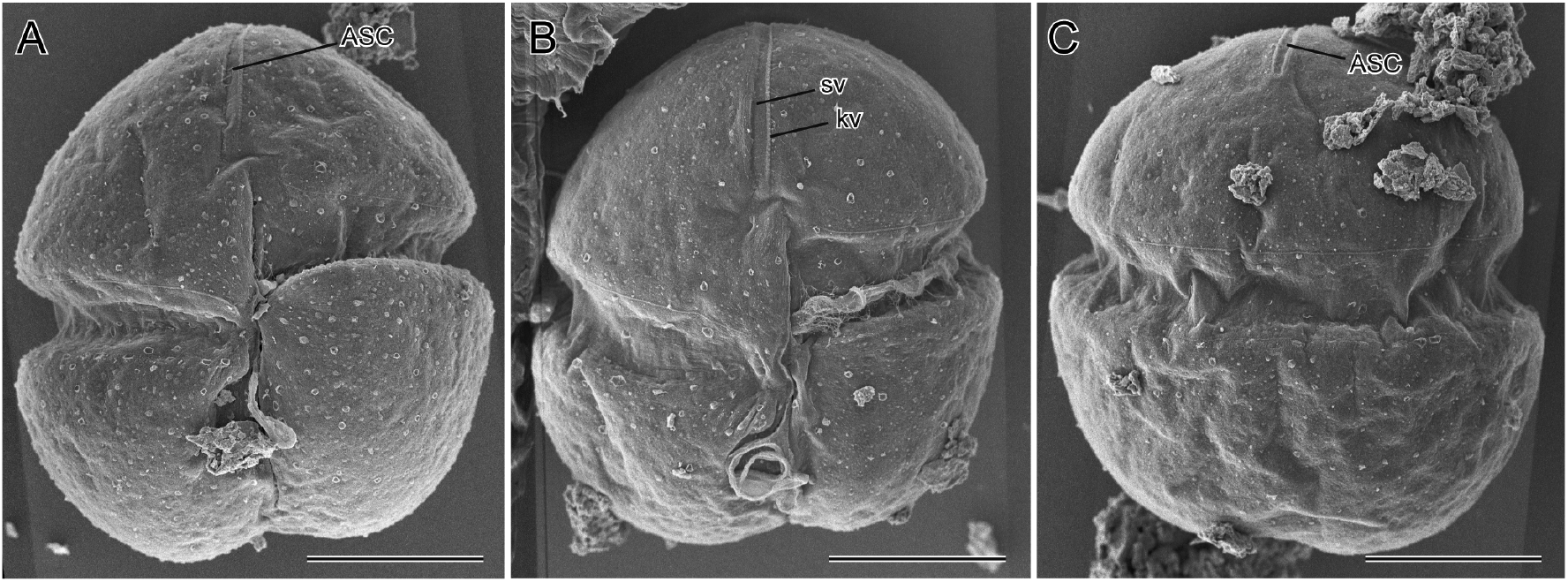
Scanning electron microscopy of *Karenia selliformis*. (A, B) Ventral views showing the straight apical structure complex (ASC, = apical groove) composed of knob vesicles (kv) and smooth vesicles (sv). (C) Dorsal view, showing the ASC and rugose cell surface. Scale bars = 10 μm.

### 3.3. Morphology of co-occurred dinoflagellates of the Kareniaceae

Six other kareniacean dinoflagellate species were found in the *Kr. selliformis* bloom; *Karenia longicanalis*, *Kr. mikimotoi*, *Karlodinium* sp., *Takayama* cf. *acrotrocha*, *Takayama tuberculata* de Salas, and *Takayama* sp. (Fig. 4).

**Fig. 4.**
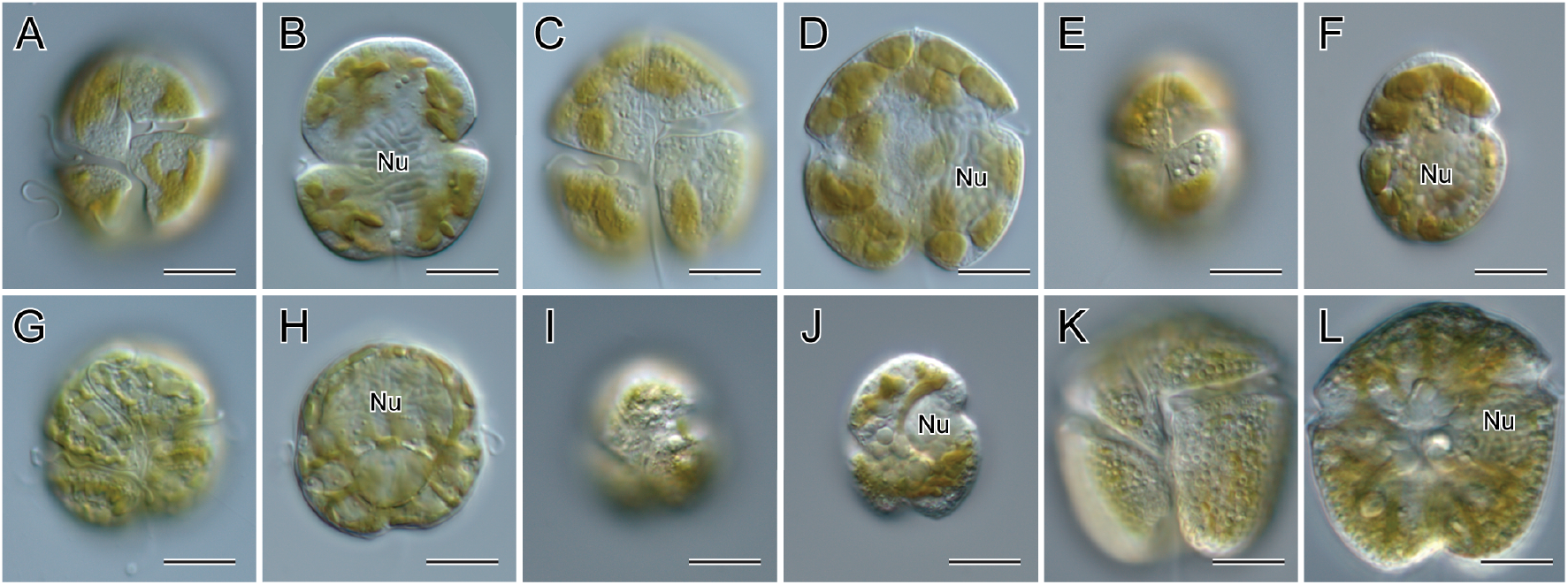
Cells of kareniacean dinoflagellates found in the bloom of *Karenia selliformis*, light microscopy. (A, B) *Karenia longicanalis*. (C, D) *Karenia mikimotoi*. (E, F) *Karlodinium* sp. (G, H) *Takayama* cf. *acrotrocha*. (I, J) *Takayama tuberculata*. (K, L) *Takayama* sp. Positions of nucleus (Nu) were shown. Scale bars = 10 μm.

Cells of *Kr. longicanalis* were ovoid, 28.6–37.6 μm (mean 32.9±2.2 μm, *n* = 40) long and 23.4–30.7 μm (mean 26.5±2.0 μm, *n* = 40) wide in culture condition, with the epicone and hypocone similar in size, or a slightly wider hypocone (Fig. 4A, B). The cingulum displaced *ca*. three times of its own width. The straight ASC was long, and the lowermost part in the ventral was bending to the right. Striations could be observed on the epicone (Fig. 4A). The nucleus was ellipsoid and positioned at the center or slightly posterior in the cell (Fig. 4B). Brownish chloroplasts, with a central pyrenoid and several chloroplast lobes radially extended, were located peripherally.

Cells of *Kr. mikimotoi* were dorsoventrally flattened, 24.6–35.1 μm (mean 31.2±3.0 μm, *n* = 30) long and 21.9–30.9 μm (mean 26.8±2.4 μm, *n* = 30) wide in culture condition, with the roundish conical epicone and hemispherical hypocone with an antapical depression (Fig. 4C, D). The ASC was straight. The nucleus was elliptical and positioned in the left side, or spherical in the left side of the hypocone (Fig. 4D). Brownish and bean-shaped chloroplasts, each containing a wedge-shaped pyrenoid, were located peripherally, mainly in the dorsal side of the cell.

Cells of *Karlodinium* sp. were ellipsoid, with the epicone slightly smaller than the hypocone (Fig. 4E, F). The cingulum positioned at median, or slightly anterior, with the displacement of its cingular width. The ASC was straight and started from right side of the anterior sulcal extension (Fig. 4E). The nucleus was spherical or elliptical and positioned in the posterior part of the cell (Fig. 4F). Chloroplasts were ovoid and positioned in the periphery of the cell. This species was rare in the bloom samples and the culture was not established.

Cells of *Takayama* cf. *acrotrocha* were spherical, 23.5–31.4 μm (mean 26.8±1.8 μm, *n* = 30) long and 20.7–28.2 μm (mean 24.5±1.7 μm, *n* = 30) wide in culture condition, and the epicone was smaller than the hypocone (Fig. 4G, H). The ASC was deep and sigmoid in shape (Fig. 4G). The nucleus was laterally elongated, and anteriorly positioned in the dorsal side. A large spherical pyrenoid was conspicuous, which was positioned in the center of the hypocone (Fig. 4H). Brownish chloroplasts were radially extended from the pyrenoid, then irregularly reticulated in the periphery.

Cells of *Takayama tuberculata* were ellipsoid, with the conical epicone smaller than the hypocone, and had an antapical excavation (Fig. 4I, J). The cingular displacement was large, and the ASC was sigmoid (Fig. 4I). The spherical nucleus was positioned in the dorsal side (Fig. 4J). Compound pyrenoid, each spherical and about ten in number, was positioned in the center or right side of the hypocone (Fig. 4J). Chloroplasts were extended from the pyrenoid. This species was rare in the bloom samples and the culture was not established.

Cells of *Takayama* sp. were large, 35.2–48.8 μm (mean 41.9±4.2 μm, *n* = 18) long and 33.0–47.6 μm (mean 40.1±4.4 μm, *n* = 18) wide in the bloom samples, with the conical epicone smaller than the hypocone (Fig. 4K, L). The hypocone was bilobed with the antapical excavation. The surface of the hypocone was reticulated, particularly in the posterior part of the cell (Fig. 4K). The ASC was sigmoid and deep (Fig. 4K). The nucleus was laterally elongated at the dorsal, and positioned near the cingulum (Fig. 4L). Compound pyrenoid was positioned in the center of the cell, consisted of spherical pyrenoids of similar size. Chloroplasts were extended from the spherical pyrenoid to the periphery.

### 3.4. *Chloroplasts of* Karenia longicanalis, Kr. mikimotoi *and* Kr. selliformis *in the bloom*

For an unambiguous identification of *Karenia* species in the bloom, chloroplast morphology of *Kr. longicanalis*, *Kr. mikimotoi* and *Kr. selliformis* were compared (Fig. 5, Table 2). Chloroplasts of *Kr. longicanalis* were multilobed, with lobes irregularly extended from the central pyrenoid (Fig. 5A, B). They ranged 9.0–19.4 μm (mean 14.5±2.4 μm, *n* = 50) in major axis, and 4–7 (mean 6.1±0.6, *n* = 40) in number in culture condition. Chloroplasts of *Kr. mikimotoi* were ellipsoid or bean-shaped, with a wedge-shaped pyrenoid laterally located (Fig. 5C, D). They ranged 4.7–10.6 μm (mean 7.3±1.5 μm, *n* = 50) in major axis, and 13–39 (mean 24.5±6.4, *n* = 40) in number in culture condition. Chloroplasts of *Kr. selliformis* were variable in shape, from subspherical to strap-shaped, irregularly elongated and curved, but not branched, with a pyrenoid (Fig. 5E, F). The subspherical forms were small, ranged 2.9–4.6 μm (mean 3.7±0.4 μm, *n* = 50) in diameter, and the strap forms ranged 6.2–15.1 μm (mean 9.4±1.9 μm, *n* = 50) in major axis. Chloroplasts were numerous in a cell of *Kr. selliformis*, counted 46–105 (mean 70.6±15.8, *n* = 30) in number in culture condition.

**Fig. 5.**
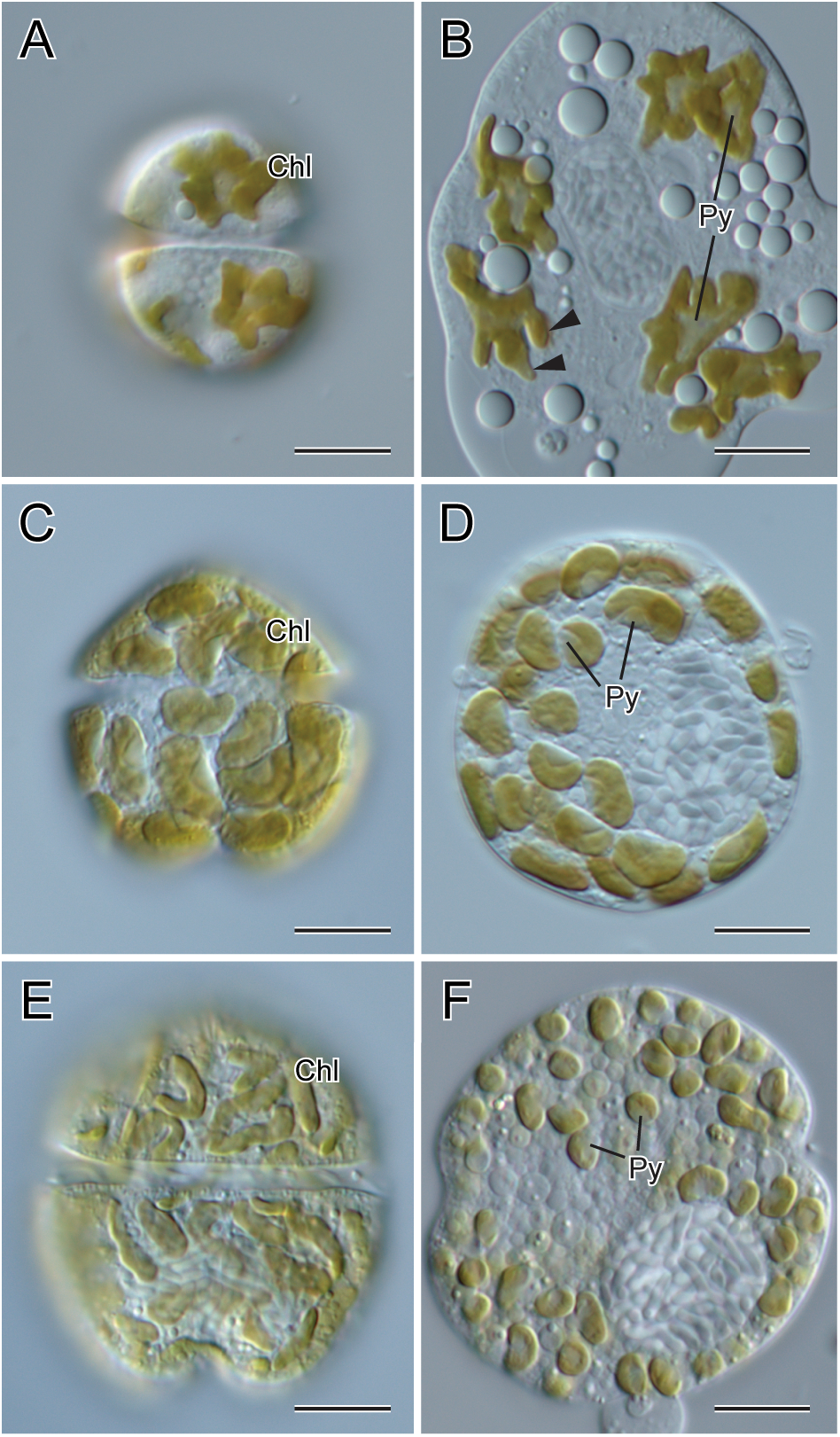
Chloroplast morphology of *Karenia longicanalis* (A, B), *Kr. mikimotoi* (C, D), and *Kr. selliformis* (E, F), note the differences in shape, size, and number. Dorsal views of motile cells (A, C, E) and cells squashed (B, D, F), showing chloroplasts (Chl), pyrenoids (Py) and chloroplast lobes of *Kr. longicanalis* (arrowheads). Scale bars = 10 μm.

### 3.5. Phylogeny of kareniaceans in the bloom Hokkaido 2021

ITS and LSU rDNA sequences determined from kareniacean dinoflagellates in this study were five cultures of *Kr. longicanalis*, four cultures of *Kr. mikimotoi*, four cultures and 15 field isolates of *Kr. selliformis* (Supplementary Fig. S2), a culture of *Takayama* cf. *acrotrocha*, and three cultures of *Takayama* sp. (Table 1).

In the ITS phylogeny, *Karenia* species formed a clade (Bayesian posterior probabilities/ML bootstrap support = 1.00/100%), which contained clades of *Kr. longicanalis* (1.00/99%), *Kr. selliformis* (1.00/99%), and other *Karenia* species (Fig. 6). The clade of *Kr. longicanalis* was subdivided into three groups; the group I consisted of isolates from China and Korea (1.00/87%), group II consisted cultures from New Zealand and Aomori, Japan (1.00/99%), and group III consisted only of four cultures from Hokkaido, Japan. The clade of *Kr. selliformis* (1.00/99%) was divided into the group I and II, which has previously been reported in Mardones et al. (2020). The group I contained cultures/isolates from Chile, Russia, and Hokkaido, Japan (0.99/74%). The group II consisted of cultures from New Zealand, Iran, Tunisia and Aomori, Japan (1.00/97%). All ITS sequences of cultures/isolates from Hokkaido, Japan and Kamchatka, Russia were identical.

**Fig. 6.**
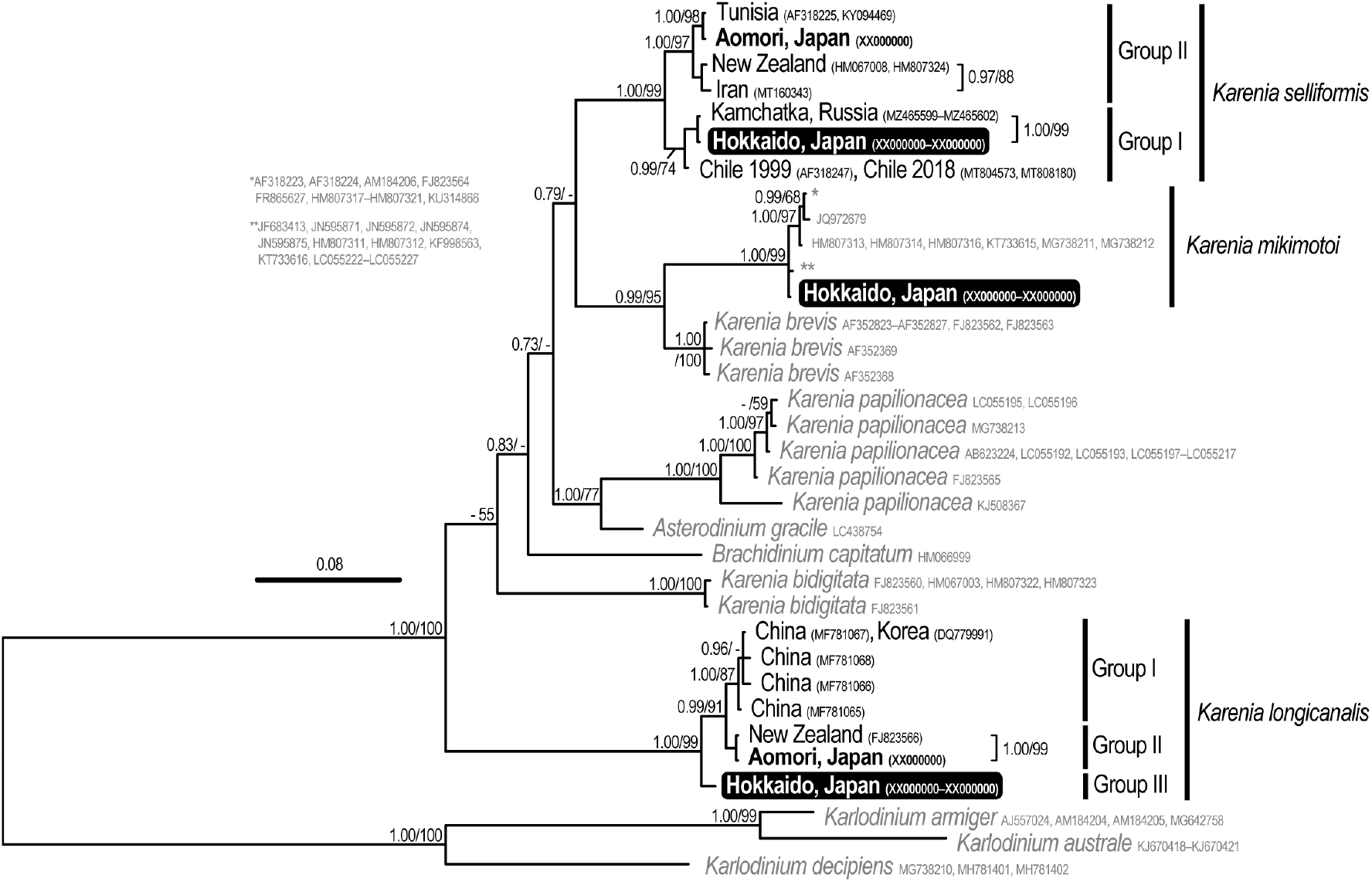
Bayesian phylogeny inferred from ITS region (34 OTUs, 720 characters); posterior probabilities (PP) and bootstrap support (BS) values of ML indicated; support values of ≥ 0.7 for PP and ≥ 50% for BS shown; DNA sequences analysed in this study are highlighted by boldface. Sequences from the Hokkaido bloom are highlighted in black box, and other sequences determined in this study were indicated in boldface.

In the LSU rDNA phylogeny, species of *Karenia* formed a clade (1.00/99%), which contained clades of *Kr. longicanalis* (1.00/98%), *Kr. mikimotoi* (1.00/99%), *Kr. selliformis* (1.00/94%) and other *Karenia* species (Fig. 7). In the clade of *Kr. longicanalis*, three sub-clades, group I, II, and III, were found. The group I composed of isolates from China, Hong Kong, Korea, South China Sea, and Russia (0.98/66%). The group II consisted of cultures form Australia, Canada, France, New Zealand and Aomori, Japan (0.92/51%). The group III included the cultures from France and Hokkaido, Japan (0.97/91%). *Karenia mikimotoi* cultures from Hokkaido had identical sequences, which branched in the *Kr. mikimotoi* clade (1.00/99%), and the relationship in the *Kr. mikimotoi* clade was unclear. In the *Kr. selliformis* clade (1.00/94%), members were separated into two sub-clades, group I and II. The group I consisted of isolates from Chile 1999, Russia, and Hokkaido, Japan (0.84/64%). All isolates of *Kr. selliformis* from Hokkaido, Japan and Kamchatka, Russia had identical LSU rDNA sequences. The group II composed of cultures/isolates from China, Chile 2018, New Zealand, Tunisia and Aomori, Japan (1.00/72%). This group II encompassed the phylotypes II and III in Mardones et al. (2020). Species of *Takayama* formed a clade (1.00/99%), which was related to the clade of *Karlodinium* species. A culture of *T.* cf. *acrotrocha* branched in the clade of *T. acrotrocha/Takayama xiamenensis* Gu*/Takayama helix* de Salas, Bolch, Botes et Hallegraeff (0.98/74%), and four cultures/isolates *Takayama* sp. were positioned in the clade of *Takayama tasmanica* de Salas, Bolch et Hallegraeff*/T. tuberculata* (1.00/94%).

**Fig. 7.**
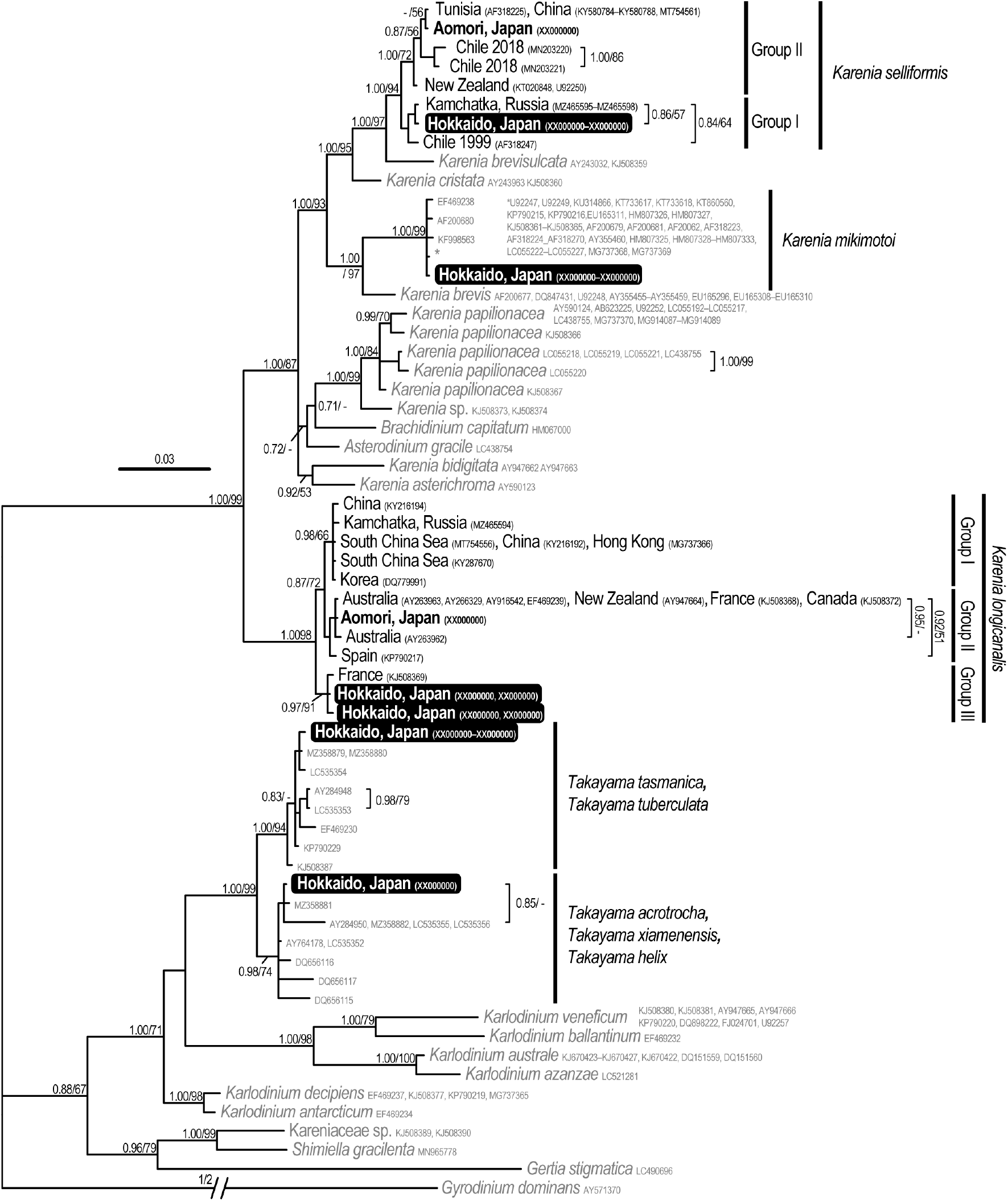
Bayesian phylogeny inferred from partial LSU rDNA (D1–D3) sequences (63 OTUs, 972 characters); posterior probabilities (PP) and bootstrap support (BS) values of ML indicated; support values of ≥ 0.7 for PP and ≥ 50% for BS shown; DNA sequences analysed in this study are highlighted by boldface. Sequences from the Hokkaido bloom are highlighted in black box, and other sequences determined in this study were indicated in boldface.

## 4. Discussion

### 4.1. *Blooms of* Karenia selliformis *and associated fisheries damage*

The dominant species of the bloom along the Pacific coast of eastern Hokkaido, Japan in September–November 2021 was identified to *Kr. selliformis* based on its morphological and molecular characteristics. This is the first blooming case of *Kr. selliformis* in Japan, although a single cell was previously isolated and observed from Hiranai, Aomori in 2019 (Benico, 2020; this study).

So far, cells of *Kr. selliformis* have been found from various regions, such as the Southwest Pacific (Australia and New Zealand), Southeast Pacific (Chile), Northwest Pacific (Japan and Russia), Northeast Pacific (Mexico), Arabian Gulf (Iran, Kuwait and Saudi Arabia), Mediterranean (France and Tunisia), and Northwest Atlantic (U.S.) (MacKenzie et al., 1996; Clément et al., 2001; Heil et al., 2001; Uribe and Ruiz, 2001; Guillou et al., 2002; Haywood et al., 2004, 2007; Hallegraeff et al., 2010, 2021b; Feki et al., 2013; Escobar-Morales and Hernandez-Becerril, 2015; Mardones and Clément, 2016; Prabowo and Agusti, 2019; Benico, 2020; Harris et al., 2020; Mardones et al., 2020; Anderson et al., 2021; Sunesen et al., 2021; Zingone et al., 2021). In early 1993, *Gymnodinium* sp., possessing the laterally elongated nucleus in the hypocone, was first reported from the North Island, New Zealand (fig. 1 in Chang, 1995; fig. 2 in MacKenzie et al., 1995). Subsequently, this species bloomed in the coasts of the South Island in 1994, and mortality of a variety of marine organisms was recorded (MacKenzie et al., 1996). *Karenia selliformis* was originally described in Haywood et al. (2004), using the culture CARD79 isolated from the Foveaux Strait, South Island, New Zealand in 1994. Its negative impacts to marine fauna have been documented in several regions. Blooms of *Kr. selliformis* have been found in relatively warmer locations associated with fish kills; in the Gulf of Gabes, Tunisia in 1994 (Feki et al., 2013; Zingone et al., 2021), and in Kuwait and adjacent locations in the Arabian Gulf in September–October 1999 (Heil et al., 2001). In addition, in 1999, a massive bloom of the species caused mortalities of salmon, sea urchins, limpets, scallops, etc., in Punta Arenas, Chile (Clément et al., 2001; Uribe and Ruiz, 2001; Mardones and Clément, 2016), and it recurred in the western Patagonian coast in 2018, associated with mortalities of farmed salmon, sea urchins, octopus, sea stars and several species of bivalves (Mardones et al., 2020). In September–October 2020, an intensive bloom caused severe mortalities of marine organisms along the coast of Kamchatka, Russia (Orlova et al., in preparation), and *Kr. selliformis* was detected from the bloom.

Among these records of *Kr. selliformis* bloom, the Hokkaido bloom 2021 resembles those in Chile and Russia in terms of the lower blooming temperature and traits of dead organisms. The seawater temperature ranged 9.8–17.6°C (records of >100 cells mL^−1^) in Hokkaido is comparable to the bloom in Chile in 1999; higher cell density was recorded under the temperature over 13.5°C (Uribe and Ruiz, 2001). In Kamchatka, Russia in 2020, the sea surface temperature range was approximately 9–13°C from satellite image (Bondur et al., 2021), when the cells of *Kr. selliformis* were detected from the bloom (Orlova et al., in preparation). During the blooms of *Kr. selliformis* in Chile, Japan and Russia, mortalities of fish, sea urchins, octopus, and other benthic invertebrates were found (Mardones et al., 2020; Orlova et al., in preparation). From *Kr. selliformis*, bioactive compounds such as gymnodimine and brevenal-related compounds have also been detected (e.g., Seki et al., 1995; MacKenzie et al., 1996; Mardones et al., 2020). Therefore, lethal effects to marine organisms and producing compounds would be important topics for *Kr. selliformis* in the Hokkaido bloom 2021.

### 4.2. *Identification of* Karenia selliformis *and other kareniaceans in the bloom*

Difficulty in light microscopic identification of kareniacean species attributes to their morphological plasticity and co-occurrence of other kareniaceans in a bloom. In the previous *Kr. selliformis* blooms, two different morphotypes have been documented (e.g., Uribe and Ruiz, 2001). Co-occurrences of several kareniaceans have so far been observed in a bloom, e.g., *Kr. brevis* bloom in the Gulf of Mexico, U.S., and *Karenia* spp. bloom in the Gulf of Penas, Chile (Heil and Steidinger, 2009; Villanueva et al., 2017). To clarify the morphological variability of *Kr. selliformis* in the Hokkaido bloom, we attempted single cell PCR and culture establishment for different cell shapes of *Kr. selliformis*, followed by identifications of other kareniaceans present in the bloom. All results of molecular identification and culture establishment showed *Kr. selliformis* as the dominant species, and its identical rDNA sequences allowed us to elucidate the morphological variation of *Kr. selliformis* in the bloom.

Diagnostic characters of *Kr. selliformis* in the Hokkaido bloom, by which it could be distinguished from other *Karenia* species such as *Kr. longicanalis* and *Kr. mikimotoi*, are the posteriorly located nucleus and numerous small-sized chloroplasts. The laterally elongated nucleus located in the hypocone corresponds to previous reports of *Kr. selliformis* (e.g., Haywood et al., 2004; Mardones et al., 2020). The nucleus was spherical in the smaller, ellipsoidal, or transparent cells, but it was consistently positioned in the hypocone. On the other hand, as for the chloroplasts, its size and shape are variable in a species and even in a strain, and generally not used as a reliable diagnostic character in dinoflagellates. Likewise for the chloroplast in *Kr. selliformis*, the size has not been compared with other *Karenia* species previously, and the different numbers have been reported in *Kr. selliformis* cells; several chloroplasts were found in the cell from New Zealand in 1994 and Chile in 2018 (Haywood et al., 2004; Mardones et al., 2020), 18–20 chloroplasts from New Zealand in 1994 (MacKenzie et al., 1996), and 56±9 (*Gymnodinium* sp.1) and 34±8 (*Gymnodinium* sp.2) chloroplasts from Chile bloom in 1999 (Uribe and Ruiz, 2001). In Hokkaido bloom 2021, about 70 chloroplasts were observed in typical cells of *Kr. selliformis*. Although the shape was variable from granular subspherical to strap-like elongated form, the sizes of the subspherical form were approximately 3.7 μm in diameter, which were apparently small and discernable from those in *Kr. longicanalis* and *Kr. mikimotoi*. The numerous and granular chloroplasts helped identification of *Kr. selliformis* in the bloom, especially for its smaller and ellipsoidal cell forms.

Cell sizes and morphology of *Kr. selliformis* observed in the Hokkaido bloom 2021 were larger than those in previous reports (Table 2). The size range of typical cells in the Hokkaido bloom, 35.3–43.6 μm (mean 39.4 μm) in length, is larger than those reported from New Zealand in 1994 (26–32 μm, MacKenzie et al., 1996; 20–27 μm, Haywood et al., 2004), Kuwait in 1999 (17–37 μm, Heil et al., 2001), Mexico (22–24 μm, Escobar-Morales and Hernandez-Becerril, 2015), and Chile in 2018 (25–30 μm, Mardones et al., 2020), but similar to cells found in Punta Arenas, Chile in 1999 (38–48 μm of *Gymnodinium* sp.1 and 32–48 μm of *Gymnodinium* sp.2, Uribe and Ruiz, 2001).

The differences in the cell size and chloroplast number are probably due to its morphological plasticity and the variability under culture condition. Cell sizes and chloroplast numbers were measured and counted directly from cells in the blooms of Chile in 1999, Kuwait in 1999 and Hokkaido, Japan in this study (Heil et al., 2001; Uribe and Ruiz, 2001), while they were obtained from cultured cells of the blooms of New Zealand in 1994 and Chile in 2018 (MacKenzie et al., 1996; Haywood et al., 2004; Mardones et al., 2020). The other possibility is their phylogeny, the group I and II, in *Kr. selliformis*. The larger cells about 40 μm in length with numerous chloroplasts were observed from the blooms of Chile in 1999 and Hokkaido, Japan in 2021; both belong to the group I of *Kr. selliformis*. To clarify the morphological differences between the group I and II, further observation of blooming cells of *Kr. selliformis* (group II) is required.

### 4.3. *Transparent cells of* Karenia selliformis

Transparent motile cells of *Kr. selliformis*, with a small number of reduced chloroplasts or lacking, were first recognized in the dense *Kr. selliformis* bloom in early October, under the seawater temperature of approximately 13–14°C, and were observed until the end of the bloom in November, under *ca*. 8–9°C. Accumulation of larger oil droplets ca. four in number is also a feature of this transparent cell. Motile cells lacking chloroplasts are unusual in the Kareniaceae, although resting cysts of *Kr. mikimotoi* show a similar appearance in the cytoplasm (Liu et al., 2020). According to Liu et al. (2020), resting cysts of *Kr. mikimotoi* are formed from the planozygote, with two longitudinal flagella, that finally become a spherical immotile cell with shrunken chloroplasts. Oil droplet-like structures, resembling those in the *Kr. selliformis* transparent cells, are also seen (fig. 1k and m in Liu et al., 2020). It is interesting if the transparent cells of *Kr. selliformis* are planozygote or planomeiocyte of the resting cyst, which can survive in the low temperature. However, all observed transparent cells were motile and possessing only a longitudinal flagellum, which are different from the planozygote or planomeiocyte of dinoflagellates.

### 4.4. *Population of* Karenia selliformis *in the bloom*

In the Northwest Pacific, including East and Southeast Asia, cells of *Kr. selliformis* have been found in Japan (Aomori) and Russia (Benico, 2020; Orlova et al., in preparation). Moreover, metabarcoding environmental DNAs have shown the presence of *Kr. selliformis* in China and Gulf of Thailand (Fu et al., 2021). This may suggest the wide distribution of *Kr. selliformis* in this region, even at low cell density that resulted in scarce observations of the cells. These *Kr. selliformis* from China, Japan (Aomori) and the Gulf of Thailand belong to the intraspecific group II of *Kr. selliformis* (Fu et al., 2021; this study). The bioactive compounds, gymnodimines, have been detected from *Kr. selliformis* (group II) in New Zealand and Tunisia (Seki et al., 1995; MacKenzie et al., 1996; Dragunow et al., 2005; Ben Naila et al., 2012). Therefore, detections of the compounds from the South China Sea, East China Sea and Bohai Sea may also imply the distribution of *Kr. selliformis* (group II) in the Northwest Pacific, but further confirmation is needed because it can also be produced by *Alexandrium ostenfeldii* (Paulsen) Balech et Tangen (He et al., 2019; Liu et al., 2019; Ji et al., 2022). On the other hand, cells of *Kr. selliformis* proliferated in Hokkaido in September–November 2021 had identical ITS-LSU rDNA sequences to those from Kamchatka in September–October 2020, which belong to *Kr. selliformis* (group I). The molecular data suggest that the population found in Hokkaido and Kamchatka differs from the *Kr. selliformis* (group II) and is probably distributed in the Northwest Pacific. Furthermore, cells possessing numerous granular chloroplasts, and transparent cells with the straight ASC and oil droplets, were observed also in Kamchatka bloom in 2020 (Orlova et al., in preparation). According to Kuroda et al. (2021), the increase of satellite-derived chlorophyll concentration was first detected at the beginning of September 2021, in the south of Etorofu Island, and subsequently expanded south-westward to the Pacific coast of eastern Hokkaido. If the cells of *Kr. selliformis* in Hokkaido bloom had been transported from the Kamchatka Peninsula, this seed population had to survive during winter in the coastal Oyashio and Oyashio waters with the water temperature below 2°C even around 100 m in depth (Isada et al., 2019, 2021). Overwintering strategy of *Kr. selliformis* is therefore important and should be determined to understand the blooming mechanism of this marine fauna-destructive species. Regarding other kareniacean dinoflagellates found in the Hokkaido bloom 2021, e.g., *Kr. longicanalis*, the genotypes of Hokkaido and Kamchatka blooms were different; strains of Hokkaido and Kamchatka belonged to group I and group II, respectively. Associated kareniacean dinoflagellates are often found in *Karenia* blooms, but the composition may be variable in population level.

The bloom of *Kr. selliformis* along the Pacific coasts of eastern Hokkaido, accountable for mass mortality of marine organisms, had wide societal impacts in Japan. Questions on HABs are best addressed on a species-by-species and site-by-site basis, and consideration on the respective impacts on local human activities should be made (Hallegraeff et al., 2021a). This cold-water HAB caused by *Kr. selliformis* is the first case in Japan, and only several innoxious blooms have so far been recorded in eastern Hokkaido, such as *Akashiwo sanguinea* (Harasaka) Hansen and Moestrup (as *Gymnodinium splendens* Lebour) in 1983 (Kakuta, 1984; Takasugi and Kakuta, 1986). Hence, further understanding of the blooming, overwintering, and marine organism killing mechanisms of the identified species, *Kr. selliformis*, would be helpful to mitigate negative impacts of its noxious bloom.

## Acknowledgements

We thank Saho Kitatsuji, Natsuko Nakayama and Koki Yuasa of Fisheries Technology Institute, Hiroshi Shimada, Akira Miyazono, Makoto Kanamori, Masafumi Natsuike, Yutaro Ando, Akiyoshi Shinada, Masayo Nomura and crew of R/V Kinsei-maru and Hokushin-maru of Hokkaido Research Organization, for their sampling, microscopic observation and sharing bloom information. Sampling and observation were supported by members of Hokkaido Fisheries Technical Guidance Offices of Subprefectural Bureau, Hokkaido, relevant city offices, town offices, and fisheries co-operative associations. Satellite ocean color images were provided by Japan Aerospace Exploration Agency (JAXA). This work was partially supported by Japanese JSPS Kakenhi 19H03027 and 19KK0160 (MI).

---

**Supplementary Figure S1.** Video showing motile cells of *Karenia selliformis* in the bloom.

**Supplementary Figure S2.** Cells of *Karenia selliformis* used for single cell PCR.

## Notes

### Competing Interest Statement

The authors have declared no competing interest.

