## Supplementary Figure S2 for "Morphological variation and phylogeny of *Karenia selliformis* (Gymnodiniales, Dinophyceae) in an intensive cold-water algal bloom in eastern Hokkaido, Japan in September–November 2021"

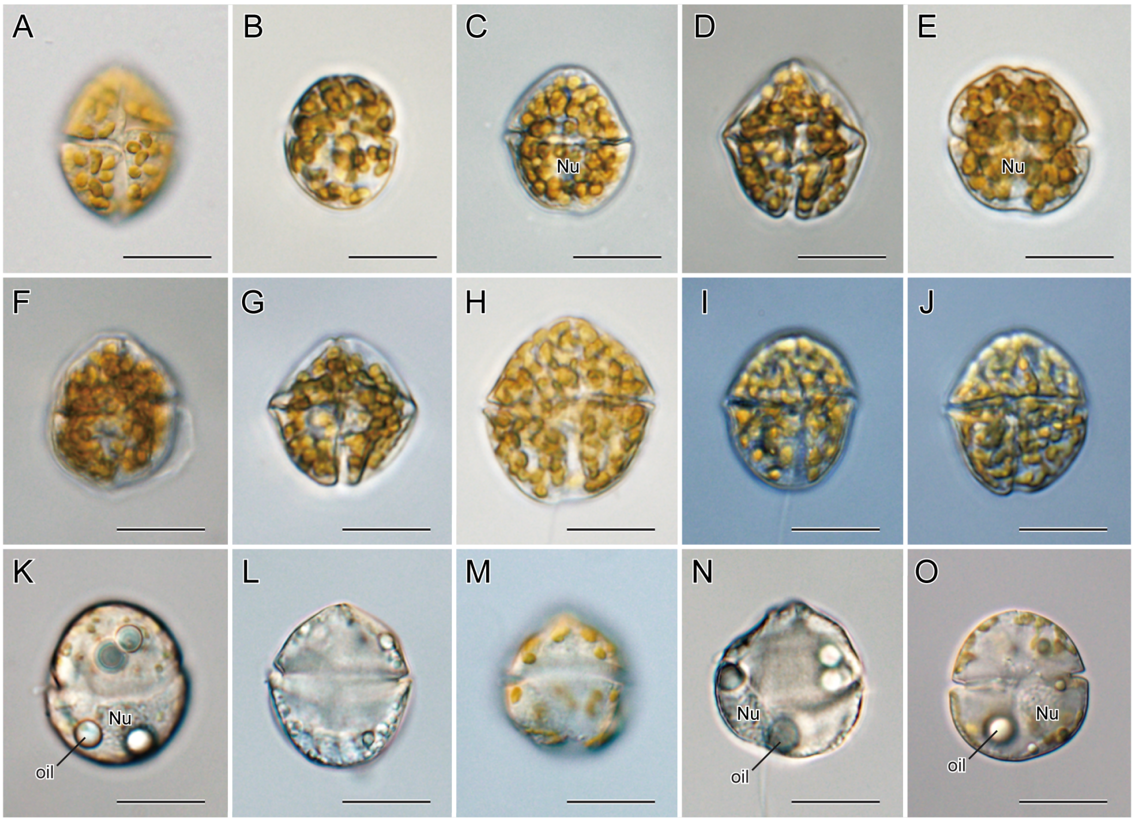


**Supplementary Figure S2.** Cells isolated for single cell PCR, light microscopy. Isolates of K1 (A), K3 (B), K7 (C), M2 (D), M3 (E), M4 (F), M5 (G), M6 (H), M7 (I), and M8 (J) from Katsurakoi Port, Kushiro, Hokkaido on 25 September 2021. Isolates with reduced chloroplast or without chloroplast, AsahiL2 (K) and AsahiL7 (L), from Asahihama Port, Taiki, Hokkaido on 13 October 2021, and OT2 (M), OT3 (N) and OT4 (O) from Otsu Port, Toyokoro, Hokkaido on 14 October 2021. Note the position of nucleus (Nu) and oil droplets (oil) in transparent cells. Scale bars = 20 µm.
